# Siderophores drive invasion dynamics in bacterial communities through their dual role as public good versus public bad

**DOI:** 10.1101/2021.06.06.447233

**Authors:** Alexandre R. T. Figueiredo, Özhan Özkaya, Rolf Kümmerli, Jos Kramer

**Affiliations:** Dept. of Quantitative Biomedicine, University of Zurich, Switzerland; Dept. of Evolutionary Biology and Environmental Studies, University of Zurich, Switzerland

**Keywords:** biological invasion, siderophores, bacterial public goods, cheating, competition, bacterial interactions, pyoverdine, *Pseudomonas*, microbial ecology, community ecology

## Abstract

Microbial invasions can compromise ecosystem services and spur dysbiosis and disease in hosts. Nevertheless, the mechanisms determining invasion outcomes often remain unclear. Here, we examine the role of iron-scavenging siderophores in driving invasions of *Pseudomonas aeruginosa* into resident communities of environmental pseudomonads. Siderophores can be “public goods” by delivering iron to individuals possessing matching receptors; but they can also be “public bads” by withholding iron from competitors lacking these receptors. Accordingly, siderophores should either promote or impede invasion, depending on their dual effects on invader and resident growth. Using supernatant feeding and invasion assays, we show that invasion success indeed decreased when the invader was inhibited (public bad) rather than stimulated (public good) by the residents’ siderophores. Conversely, invasion success often increased when the invader could use its siderophore to inhibit the residents. Our findings identify siderophores as a major driver of invasion dynamics in bacterial communities.

## Introduction

The invasion of non-native species into established communities is a widespread phenomenon and has been extensively studied in eukaryotes (Lowe *et al*. 2000), where it often leads to biodiversity loss and the deterioration of ecosystem functioning (Ehrenfeld 2010; Doherty *et al*. 2016). More recently, invasions have also received attention in microbiology because diverse microbial communities populate almost all habitats on earth (Litchman 2010; Amalfitano *et al*. 2015; Thakur *et al*. 2019), and often perform essential functions such as the fixation of CO_2_ and nitrogen or the synthesis of vitamins and amino acids (Bäckhed *et al*. 2005; Wagg *et al*. 2014; Sunagawa *et al*. 2015; Trivedi *et al*. 2020). Microbial invasions can disturb such functions, for instance by reducing the diversity or altering the composition of the resident community (Acosta *et al*., 2015; Mallon et al., 2015a), and may thereby compromise ecosystem services or spur dysbiosis and disease in hosts (Bäumler & Sperandio, 2016).

The outcome of microbial invasions is often linked to properties of the resident community and its members. High diversity and ecological dissimilarity within a community, as well as high niche overlaps between residents and invaders tend to reduce invasion success (Van Elsas *et al*., 2012; Eisenhauer *et al*., 2013; Van Nevel et al., 2013; Wei *et al*., 2015; Li *et al*., 2019). By contrast, high resource availability may either promote or hamper invasion, depending on who can better use the resources (Ma *et al*. 2015; Mallon *et al*. 2015b; Yang *et al*. 2017). Mechanistically, two broad classes of traits seem to predict whether a microorganism can successfully invade. First, metabolic versatility and high growth potentials make invaders strong competitors over limited resources (Litchman 2010; Mächler & Altermatt 2012; Ma *et al*. 2015). Second, the deployment of toxins, phages, or contact-dependent killing mechanisms allows invaders to subdue residents through interference competition (Brown *et al*. 2006; Libberton *et al*. 2015; García-Bayona & Comstock 2018).

While resource and interference competition are typically seen as distinct mechanisms acting via separate traits, we here investigate a case where both types of competition could operate through a single trait: the secretion of iron-scavenging siderophores. On the resource side, siderophores are essential for the uptake of environmentally-bound iron (Cornelis 2010). On the interference side, siderophores can be secreted beyond the level required for iron scavenging to starve competitors of iron (Niehus *et al*. 2017; Leinweber *et al*. 2018). This dual role is possible because (i) most bacterial taxa produce siderophores (Andrews *et al*. 2003); (ii) many chemically different types exist, each requiring a specific receptor for iron uptake (Cornelis & Matthijs 2002); and (iii) bacteria typically possess additional receptors for the uptake of heterologous siderophores produced by other community members (Hartney *et al*. 2011; Sexton *et al*. 2017). Secreted siderophores can therefore take on two opposing roles for community members: they can be “public goods” and promote iron acquisition and growth of members with a matching receptor; and they can be “public bads” and inhibit the growth of members lacking the matching receptor by locking iron away (Bruce *et al*. 2017; Butaite *et al*. 2017; Niehus *et al*. 2017; Gu *et al*. 2020; Kramer *et al*. 2020b).

Here, we test whether siderophores, in their role as public goods vs. public bads, can predict the success of a bacterium invading a resident community. We hypothesize that invasion success depends on the combined effect that invader and residents have on each other through their siderophores (Fig. S1). Put simply, invasion success should be lowest when the resident produces a public bad for the invader and the invader a public good for the resident. Conversely, invasion success should be highest when the resident produces a public good and the invader a public bad. Finally, invasion success should be intermediate when resident and invader both produce public bads or public goods, with the outcome probably depending on the amount of siderophores and the relative magnitude of their inhibitory or stimulatory effects. Moreover, we hypothesize that invasion success declines in more diverse resident communities because the likelihood increases that at least some of the residents’ siderophores will be public bads for the invader. Finally, we hypothesize that invasion success depends on the origin of residents, because different habitats often favor the evolution of different siderophore types and production levels (Kümmerli *et al*. 2014; Butaite *et al*. 2018).

To test our predictions, we used eight resident communities from pond and soil samples, each comprising 20 members of the genus *Pseudomonas*, and the opportunistic human pathogen *P. aeruginosa* as the invader. Invader and residents belong to the fluorescent pseudomonads that secrete pyoverdine as their primary siderophore (Butaite *et al*. 2017; Schalk *et al*. 2020). Dozens of structurally different pyoverdine types exist (Meyer et al., 2008). Each strain produces only one type, but typically has multiple receptors to take up additional pyoverdines produced by other community members (Butaite *et al*. 2017; Sexton *et al*. 2017). We first quantified growth and pyoverdine production of all 160 residents and the invader under iron-limited (pyoverdine-inducing) and iron-rich (control) conditions. Next, we fed pyoverdine-containing supernatants of all residents to the invader and vice versa. This allowed us to distinguish between pyoverdines acting as public good versus public bad, and to quantify their effects on growth. Finally, we conducted invasion-from-rare experiments, where we competed the invader against single residents or communities of 5, 10, or 20 residents.

We found that invasion success was shaped by both public good and public bad effects of pyoverdines. Specifically, invasion success increased when resident pyoverdines promoted rather than inhibited invader growth, and this effect was reinforced in competitions with single residents when the invader produced an inhibitory pyoverdine. Conversely, invasion success declined when residents produced inhibitory pyoverdines and when the invader competed against more diverse resident communities. Our findings demonstrate that siderophore-mediated interactions can have a major impact on bacterial community assembly and invasion dynamics.

## Materials & Methods

### Strain description

The residents used in our experiments were drawn from an established collection of 320 genetically characterized pseudomonads, isolated from eight soil and eight pond samples (20 isolates per sample). Details on sampling and identification procedures of these isolates are provided in Butaite *et al*. (2017; 2018). Here, we used a subset of 160 isolates from four soil and four pond samples [hereafter: community]. As the invader, we used *P. aeruginosa* PAO1 (ATCC 15692), a wound isolate that has become a laboratory model pathogen (Stover *et al*. 2000). To be able to distinguish this strain from the residents (none of which is a *P. aeruginosa* strain), we used PAO1-*mcherry*, a variant that constitutively expresses the fluorescent protein mCherry (chromosomal insertion attTn7::Ptac-*mcherry*) (Rezzoagli et al., 2019).

### Growth and pyoverdine production measurements

We quantified growth and pyoverdine production of all strains under iron-limited and iron-rich conditions. First, we grew bacterial pre-cultures in 96-well plates containing 200 µl lysogeny broth (LB) per well under shaking conditions (170 rpm) for 48h at 28°C. Subsequently, we diluted pre-cultures 100-fold into 96-well plates containing 200 µl medium per well. We used (i) iron-limited CAA medium (5g casamino acids, 1.18 g K_2_HPO_4_·3H_2_O, and 0.25 g MgSO_4_·7H_2_O per litre, supplemented with 25 mM HEPES buffer and 200 µM 2,2’-Bipyridine, a strong iron-chelator), and (ii) an iron-rich control medium, also consisting of CAA but supplemented with 40 µM FeCl_3_. We inoculated each plate with residents from one community or the invader in fourfold replication. After 24h of static incubation at 28°C, we quantified growth as optical density (OD_600_, measured at 600 nm) and pyoverdine production as relative fluorescence units (RFU_pvd_, excitation| emission at 400| 460 nm) using an Infinite M200 Pro microplate reader (Tecan Trading, Switzerland). We always obtained four measurements per well after 120 seconds of shaking.

### Supernatant assay

We explored the potential for interactions between the invader and residents via secreted products under iron-limited and iron-rich conditions by exposing the invader to supernatants collected from the residents and vice versa. We first harvested supernatants from the cultures generated in the above-described experiment by spinning them through 96-well filter plates with a 3 μm glass fiber/0.2 μm Supor membrane (AcroPrep Advance, Pall Corporation, USA) and then collecting the sterile supernatants in 96-well plates. These collection plates were sealed and stored at −20°C until further use.

Next, we diluted invader pre-cultures 100-fold into new 96-well plates and subjected them to the following three experimental conditions in four-fold replication. (i) SN_limited_: 160 µL iron-limited CAA supplemented with 40 µL resident supernatant generated under iron-limited conditions; (ii) SN_rich_: 160 µL iron-rich CAA supplemented with 40 µL resident supernatant generated under iron-rich conditions; and (iii) SN_replenished_: 160 µL iron-rich CAA supplemented with 40 µL resident supernatant generated under iron-limited conditions. As control, we used 0.85% (w/v) NaCl instead of supernatant to mimic the addition of spent medium (SN_control_). We measured invader growth [OD_600_] and overall pyoverdine production [RFU_pvd_] after 24h of incubation at 28°C under static conditions.

Subsequently, we calculated each resident’s supernatant effect on invader growth as growth effects: GE_treatment_ = (SN_treatment_ / SN_control_), where SN_treatment_ = SN_limited_, SN_rich_, or SN_replenished_, with growth values being the median supernatant effects across the four replicates. Values smaller and greater than one indicate growth inhibition and stimulation, respectively. From these measures, we calculated the net growth effect of the residents’ pyoverdines as GE_net_ = (GE_limited_ – GE_replenished_) – 1. This is possible because we used the exact same supernatants for SN_limited_ and SN_replenished_, but pyoverdines are only important for growth in the former and not in the latter condition, where iron is available in excess (Gu *et al*. 2020). Note that although GE_net_ can also reflect the effects of other siderophore types, it is highly correlated with pyoverdine production in our isolates (Fig. S2) and hence mainly reflects pyoverdine effects (see the Supplementary Material). Finally, we followed the exact same protocol to quantify the effects of invader supernatant on the growth of each resident.

### Invasion-from rare competition assays

We competed the invader against single residents and resident groups under iron-limited and iron-rich conditions. We set up competition experiments for each of our eight resident communities involving the invader and either (i) an individual resident, (ii) groups of five residents, (iii) groups of ten residents, or (iv) the entire community of 20 residents. For each community, we assembled four groups of five residents and two groups of ten residents, whereby residents were drawn at random from the pool of 20 residents without replacement. To ensure that our results are not biased by non-additive effects resulting from interactions between specific residents, we constructed a second set of five-resident and ten-resident groups by repeating the above community assembly process (Bell *et al*. 2009). This resulted in 33 different competitions per resident community (20 × [1 resident] + 2 × [4 × 5 residents] + 2 × [2 × 10 residents] + 1 × [20 residents]).

We grew pre-cultures of residents and invader from freezer stocks in 24-well plates containing 1.5 ml LB per well at 28°C under shaking conditions (170 rpm). After 48h of incubation, we washed the cells in 0.85% NaCl, measured OD_600_ of each culture (in a 1:10 dilution against a 0.85% NaCl blank), and then adjusted resident cultures to OD_600_ = 0.2 and invader cultures to OD_600_ = 0.01. Next, we assembled the resident mixes and inoculated them at a starting density of OD_600_ = 0.01 into 96-well plates containing 200 µL of iron-limited or iron-rich medium per well. Note that we used a substitutive design, whereby the overall starting density is held constant across different mixes, while individual resident density decreases when resident number increases. We then inoculated the invader at OD_600_ = 0.0001 (1% frequency) into the resident mixes. Additionally, we included invader monocultures as controls on each plate. We incubated plates at 28°C under static conditions, and measured invader growth [as RFU_mcherry_: excitation| emission at 582| 620 nm], community growth [OD_600_] and pyoverdine production [RFU_pvd_] after 16h, 20h, and 24h. Note that the constitutively expressed mCherry signal is a reliable measure of growth in competitions (Leinweber et al., 2018). We carried out competitions either in eight-fold (single residents) or twelve-fold (multi-resident communities) replication.

We calculated the invader’s success as CS_invader_ = Signal_mix_ / Signal_mono_, where Signal_mix_ and Signal_mono_ are the median mCherry signals across all replicates in the competition mix and the invader monoculture, respectively. CS_invader_ values smaller and greater than one indicate reduced and improved invader growth relative to the monoculture, respectively. In competitions with a single resident, we directly compared CS_invader_ to the pyoverdine-mediated growth effects GE_net_. In competitions with multiple residents, we compared CS_invader_ to the mean GE_net_ value across all residents in a mix.

### Statistical Analyses

We used linear mixed models (LMMs) for statistical analyses in R 4.0.3 (www.r-project.org). We implemented mixed models using the ‘lmer’ function (lme4 package: Bates *et al*. 2015), and obtained the p-values of effects in these models using the *Anova* function (car package: Fox & Weisberg 2019) and the *summary* function (lmerTest package: Kuznetsova *et al*. 2017). In a first set of LMMs, we tested whether the (square root-transformed) growth or pyoverdine production of the residents differed between experimental conditions (iron-rich vs. iron-limited) or resident habitat (soil vs. pond). The model analyzing resident growth additionally contained pyoverdine production as a covariate. In a second set of LMMs, we tested for condition-dependent and habitat-dependent differences in supernatant effects. Prior to analysis, we used the *transformTukey* function (rcompanion package: Mangiafico 2018) to find the power transformation that brought the respective supernatant effect closest to a normal distribution. In a third set of LMMs, we explored whether GE_net_ (net pyoverdine growth effect) was predictive of the invader’s success in invading resident communities. We first tested whether the (square-root transformed) success of the invader in competitions with single residents was affected by resident habitat, the effect of the invader’s pyoverdine on the resident, and/or the effect of the resident’s pyoverdine on the invader. Using the same model structure with resident number (5, 10, or 20 residents; categorical) as an additional explanatory variable, we then explored the determinants of invasion success in competitions with multiple residents. In a final LMM, we assessed whether invasion success depended on habitat, condition, resident number, or the pyoverdine production of the competition mix.

All models were initially fitted with all interaction terms. To account for the non-independence of measurements of residents from the same community, and to avoid pseudoreplication due to multiple measurements of each resident under different conditions, we initially fitted all models as random intercept models using resident community and – in case of repeated measurements – resident (mixture) identity nested within resident community as random effect(s). We then simplified models in a two-step procedure. First, we simplified the random component of the model based on likelihood ratio tests of model reduction. Second, we simplified the fixed component by dropping non-significant interaction terms (p > 0.05). Unless otherwise stated, model estimates were re-transformed to the original scale of the response for graphical display. Note that one resident hardly grew under iron-limitation, but benefitted enormously from the invader’s pyoverdine, resulting in a greatly inflated pyoverdine effect GE_net_. This caused problems in our statistical models on invasion success. Consequently, we excluded this outlier from our analysis, and set its GE_net_ = 1 for the analysis involving multi-resident communities.

## Results

### Growth and pyoverdine production profiles

We first quantified the growth of all 160 resident pseudomonads and the *P. aeruginosa* invader in monoculture (Fig. 1A). We found that growth did not differ between soil and pond residents, but was higher under iron-rich (mean ± SD: 0.799 ± 0.305) than iron-limited (0.522 ± 0.324) conditions (Table S1, Fig. 1A). The majority of residents grew worse than the invader both under iron-limited (85.5 %) and iron-rich (84.3 %) conditions.

**Figure 1.**
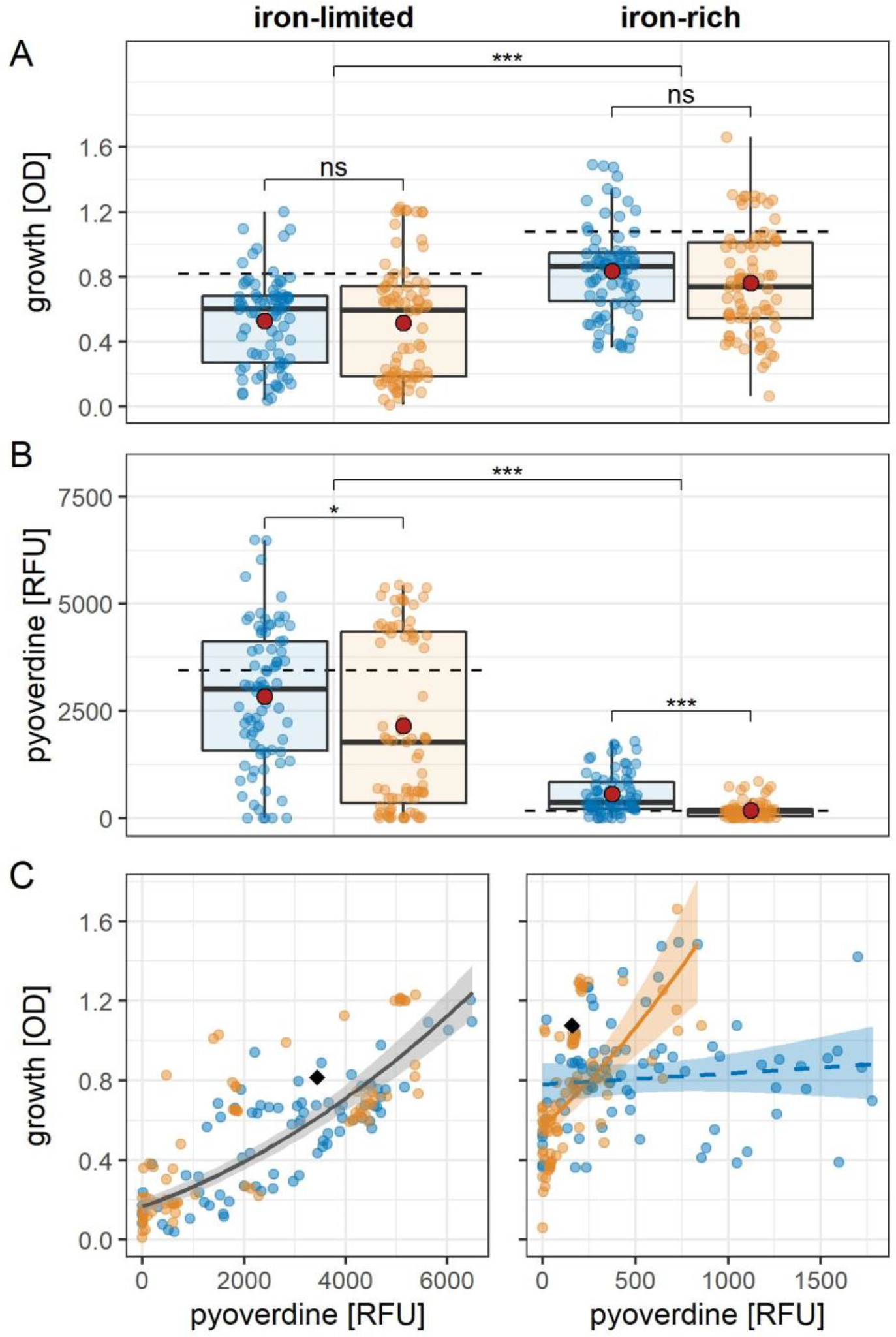
Growth and pyoverdine production of the 160 residents and the invader. Shown are the (A) growth, (B) pyoverdine production, and (C) relationship between these two measures among pond (blue) and soil (orange) residents under iron-limited and under iron-rich conditions. Boxplots show the median (bold line), mean (red point), 1st and 3rd quartile (box), and 5th and 95th percentile (whiskers). The dashed horizontal lines in (A) and (B) show the growth and pyoverdine production of the invader, respectively. The solid and dashed lines in (C) indicate significant and non-significant relationships, respectively. Shaded areas are 95% confidence intervals (grey = relationship applying to both soil and pond residents; orange = soil-specific relationship; blue = pond-specific relationship). The black diamonds in (C) show invader values for comparison.

Next, we measured the pyoverdine production of all residents and the invader. Consistent with our previous findings (Butaite *et al*. 2017; Kramer *et al*. 2020a), we observed that pyoverdine production levels varied tremendously and were significantly higher among pond than soil residents (χ^2^_1_ = 20.34, p < 0.001). Moreover, pyoverdine production was higher under iron-limited than iron-rich conditions (χ^2^_1_ = 378.35, p < 0.001; Fig. 1B), showing that residents and invader both down-regulate pyoverdine production when iron is more readily available. We further found that growth was positively linked to pyoverdine production under iron-limited conditions (t_155.80_ = 17.357, p < 0.001; Table S1, Fig. 1C), suggesting that pyoverdine production promotes growth in this environment. Under iron-rich conditions, this positive association persisted for soil residents (t_154.70_ = 6.130, p < 0.001), but vanished for pond residents (t_149.10_ = 0.825, p = 0.411; Table S1, Fig. 1C).

### The residents’ pyoverdines are predominantly public bads for the invader

To quantify to what extent resident pyoverdines serve as public goods (stimulate growth) or public bads (inhibit growth) for the invader, we fed supernatants from all 160 residents to the invader under iron-limited, iron-replenished, and iron-rich conditions, and then compared its growth relative to a control treatment. We found that the resident supernatant effects were independent of habitat (χ^2^_1_ = 0.66, p = 0.416), but differed among conditions (χ^2^_2_ = 126.79, p < 0.001). Under iron-limitation, effects varied from mild stimulation to strong inhibition (Fig. 2A). When repeating the assay with the same supernatants, but under iron-replenished conditions to cancel out pyoverdine effects, we found that most inhibitory effects were gone. Overall, supernatant effects were significantly lower (i.e. more inhibitory) under iron-limited (0.881 ± 0.265) than under iron-replenished (1.035 ± 0.106) conditions (t_316.00_ = 10.74, p < 0.001). By contrast, supernatant effects were close to neutral for both iron-replenished and iron-rich conditions (1.030 ± 0.092), suggesting a low baseline production of toxic compounds. When calculating the net pyoverdine effect on invader growth, we observed that resident pyoverdines typically had an inhibitory (public bad) effect on invader growth (Fig. 2A).

**Figure 2.**
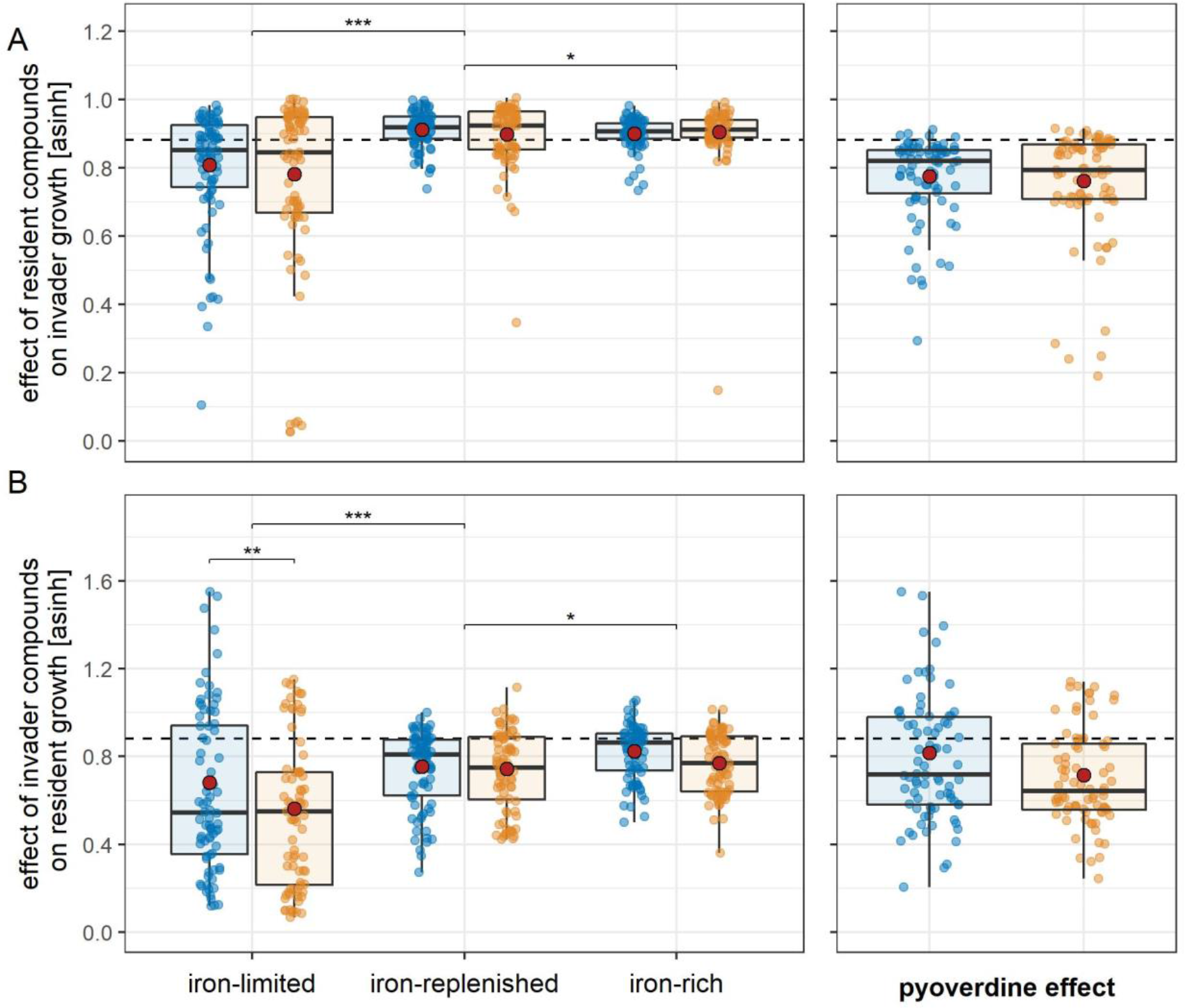
Supernatant and pyoverdine effects. Shown are (A) the effects of compounds in the resident supernatants on invader growth and (B) the effects of compounds in the invader supernatants on resident growth. Boxplots show the median (bold line), mean (red point), 1st and 3rd quartile (box), and 5th and 95th percentile (whiskers). The growth effects under iron-limitation where measured by feeding supernatant generated in iron-limited medium (supernatants contain high pyoverdine concentration + other secreted compounds) to receivers growing in iron-limited medium. Conversely, the growth effects under iron-replenished conditions where measured by feeding supernatant generated in iron-limited medium to receivers growing in iron-rich medium (such that the effect of pyoverdine is removed while the effect of other metabolites is retained). Finally, the growth effects under iron-rich conditions where measured by feeding supernatant generated in iron-rich medium (supernatants contain low pyoverdine concentration + other secreted compounds) to receivers growing in iron-rich medium. The net effect caused by pyoverdine alone (right column) was obtained by subtracting the growth effect of the iron-replenished condition from the growth effect of the iron-limited condition. The dashed horizontal lines indicate the null line where compounds in the supernatants have no effect on growth. Note that we performed an inverse hyperbolic sine (arsinh) transformation to display our data, as the high range of observed effects would have otherwise prevented a meaningful data presentation.

### The invader’s pyoverdine can be a public good or a public bad for residents

We repeated the above assay, but fed supernatants of the invader to each of the 160 residents. We found that the effect of the invader’s supernatant on resident growth depended on an interaction between condition and resident habitat (condition: χ^2^_2_ = 91.17, p < 0.001; habitat: χ^2^_1_ = 3.64, p = 0.056; interaction: χ^2^_2_ = 6.67, p = 0.036). Under iron-limitation, the supernatant effect varied between strong stimulation to strong inhibition, and was lower for soil than for pond residents (soil: 0.594 ± 0.404; pond: 0.736 ± 0.502; t_332.06_ = −2.915, p = 0.004; Fig. 2B). As before, these growth effects diminished when replenishing the supernatant with iron (iron-limited: 0.665 ± 0.460, iron-replenished: 0.821 ± 0.222, t_314.00_ = 2.953, p = 0.003), and rendered the supernatant effect similar to the effect under iron-rich conditions (0.885 ± 0.175). Calculations of the net pyoverdine effect revealed that the invader’s pyoverdine can indeed serve either as a public good or a public bad depending on the resident (Fig. 2B).

### Interactions between pyoverdine effects drive invasion success in one-to-one competitions

To test whether pyoverdine effects predict invasion success, we quantified to what extent the invader can increase in abundance from rare in competitions against each of the 160 residents. In support of our hypothesis, we found that invasion success was shaped by an interaction between the effects of the resident’s and the invader’s pyoverdines (Fig. 3A+B; Table 1A). Specifically, invasion success increased when the resident’s pyoverdine had a stimulatory (public good) rather than an inhibitory (public bad) effect on the invader, and this increase was reinforced when the invader’s own pyoverdine inhibited rather than stimulated resident growth (Fig. 3A+B). Notably, invasion success increased with the resident’s pyoverdine effect in both habitats, albeit steeper in soil (t_147.96_ = 5.531, p < 0.001) as compared to pond (t_148.71_ = 4.023, p < 0.001; Table 1A).

**Table 1.**
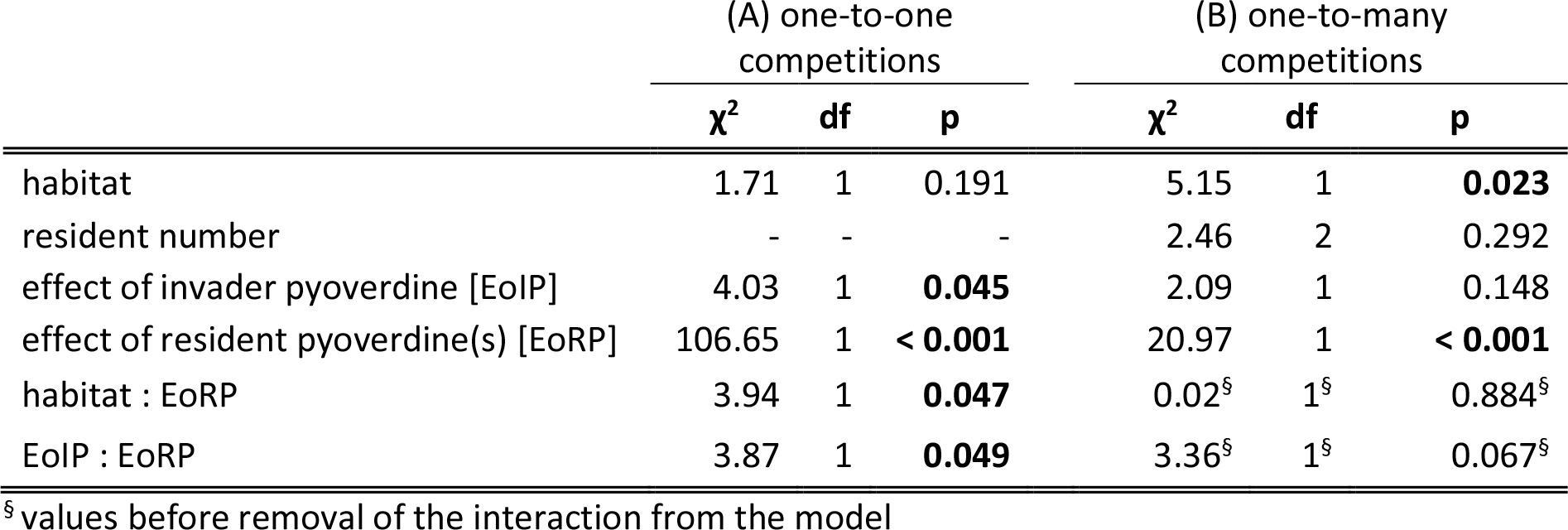
Determinants of invasion success in competitions with (A) one or (B) multiple residents. Significant p-values in bold.

|  | (A) one-to-one competitions |  |  | (B) one-to-many competitions |  |  |
| --- | --- | --- | --- | --- | --- | --- |
| | $\chi^2$ | df | p | $\chi^2$ | df | p |
| habitat | 1.71 | 1 | 0.191 | 5.15 | 1 | **0.023** |
| resident number | - | - | - | 2.46 | 2 | 0.292 |
| effect of invader pyoverdine [EoIP] | 4.03 | 1 | **0.045** | 2.09 | 1 | 0.148 |
| effect of resident pyoverdine(s) [EoRP] | 106.65 | 1 | **< 0.001** | 20.97 | 1 | **< 0.001** |
| habitat : EoRP | 3.94 | 1 | **0.047** | 0.02<sup>§</sup> | 1<sup>§</sup> | 0.884<sup>§</sup> |
| EoIP : EoRP | 3.87 | 1 | **0.049** | 3.36<sup>§</sup> | 1<sup>§</sup> | 0.067<sup>§</sup> |

<sup>§</sup> values before removal of the interaction from the model

**Figure 3.**
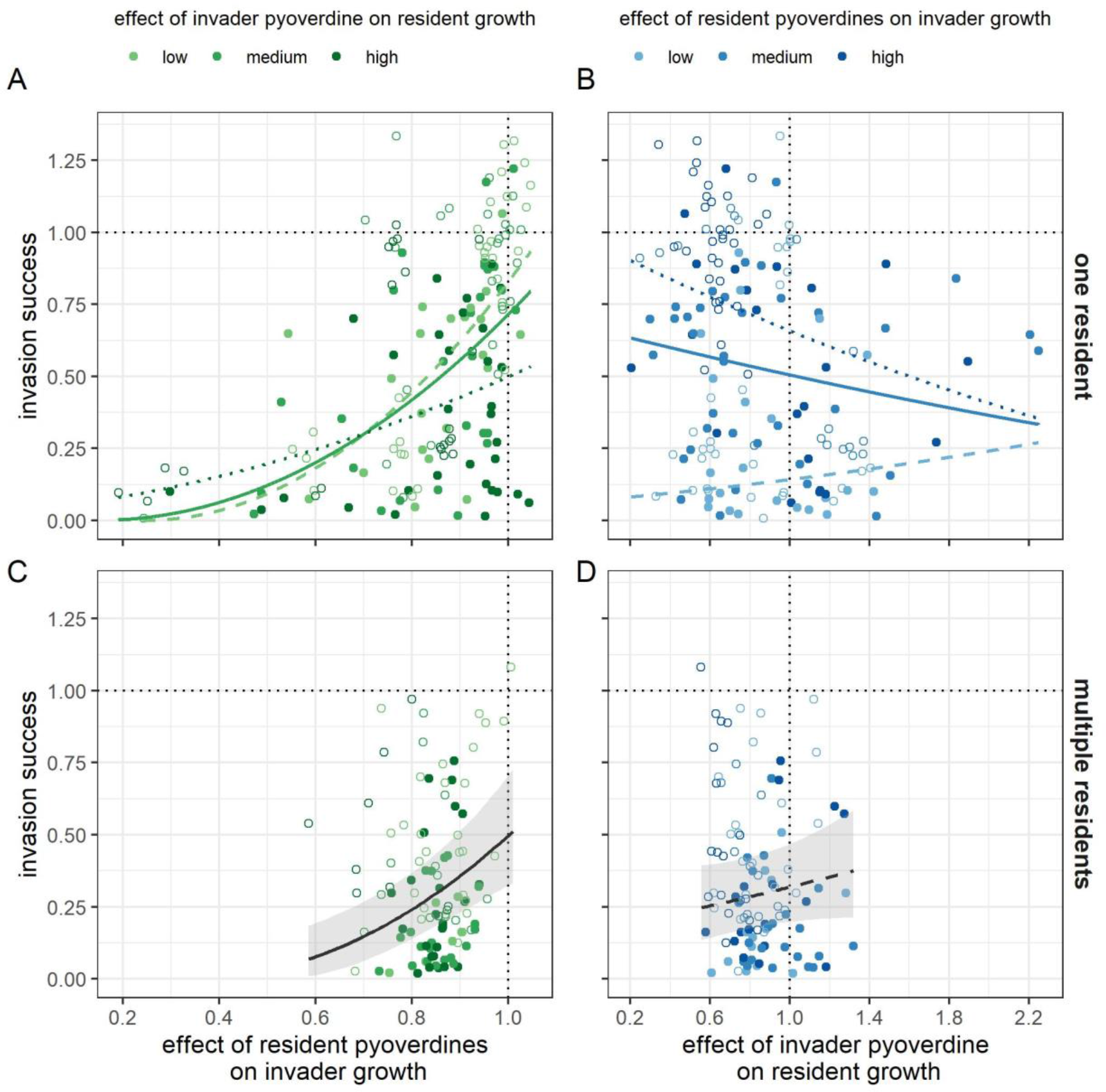
Pyoverdine-based effects on invasion success in competitions with one or multiple residents. Shown are the effects of resident pyoverdines on invader growth and the effects of invader pyoverdine on resident growth, detected in invasion-from-rare competition experiments between the invader (starting frequency 1%) and residents. (A) and (B) show the interaction between the two pyoverdine effects in competitions between the invader and a single resident with the effect of the resident pyoverdines and the effect of the invader pyoverdine as the focal predictor, respectively. (C) Main effect of the residents’ pyoverdines in competitions between the invader and multi-resident communities (5, 10, and 20 members). (D) Main effect of the invader’s pyoverdine in competitions between the invader and multi-resident communities. To illustrate the interactions in (A) and (B), regression lines are provided for a low (dashed line; mean of first tercile), medium (solid line; mean of second tercile), and high (dotted line; mean of third tercile) value of the effect of invader pyoverdine on resident growth and the effect of resident pyoverdines on invader growth, respectively. The solid and dashed regression lines in (C) and (D) indicate significant and non-significant relationships, respectively. Grey shaded areas are 95% confidence intervals. In all panels, data points are colored according to which tercile they fall in. Filled and empty points represent pond and soil residents, respectively. The dotted horizontal and vertical lines are reference lines showing invader growth in the absence of a competitor (y-axis) and invader (A+C) and resident (B+D) growth in the absence of supernatants.

### The average effect of resident pyoverdines predicts invasion success in complex communities

We then quantified invasion success into multi-resident communities, where the invader has to cope with many residents producing a cocktail of different pyoverdines. Here, we expected the effect of the invader’s pyoverdine to diminish and invasion success to be predominantly determined by the aggregate effect of the residents’ pyoverdines. Our analysis supports this prediction (Fig. 3): invasion success increased when the average effect of the residents’ pyoverdines on the invader was more stimulatory (public good) than inhibitory (public bad) (Table 1B; Fig. 3C). By contrast, the average effect of the invader’s pyoverdine on the residents had no longer an impact on invasion success, and there was no longer an interaction between the effects of the residents’ and invader pyoverdines (Fig. 3D; Table 1B). Notably, all these effects were independent of resident number (Table 1B).

### Invasion success is habitat-specific and declines in more diverse resident communities

While the above analyses directly related invasion success to public good vs. public bad properties of pyoverdines under iron-limited conditions, we now explore the effects of resident number, iron availability, habitat type, and overall community pyoverdine production on invasion success (Fig. 4). We found that invasion success was shaped by all these factors (Table 2). In particular, the invader’s success was higher in competitions with single residents (0.48 ± 0.36) than in competition with groups of five (0.27 ± 0.25), ten (0.25 ± 0.24), or twenty (0.18 ± 0.15) residents (single vs. five: t_499.5_ = −2.721, p = 0.007; five vs. ten: t_492.1_ = −0.121, p = 0.904; ten vs. twenty: t_483.2_ = −0.545, p = 0.586; Fig. 4A). This diversity effect occurred in both iron-limited and iron-rich conditions and independently of resident habitat. Overall, invasion success was higher under iron-limited as compared to iron-rich conditions (0.47 ± 0.36 vs. 0.31 ± 0.30, Table 3, Fig. 4B). Moreover, while invasion success did not vary between habitats under iron-rich conditions (t_9.74_ = 1.524, p = 0.159), the invader was more successful in competitions against soil as opposed to pond residents under iron-limited conditions (t_13.51_ = 3.586, p = 0.003; Fig. 4B). Finally, invasion success decreased with the overall pyoverdine production of the mixed culture both under iron-limited (habitat independent: t_235.3_ = −6.786, p < 0.001; Fig. 4C) and iron-rich conditions (habitat dependent, soil: t_252.9_ = −4.929, p < 0.001; pond: t_255.5_ = −4.474, p < 0.001; Fig. 4D). These results suggest that abiotic factors, such as media composition and habitat type, influence invasion success in addition to socio-ecological factors involving pyoverdine production and resident number.

**Table 2.** General determinants of invasion success. Significant p-values in bold.

| | $\chi^2$ | df | p |
| --- | --- | --- | --- |
| habitat | 0.89 | 1 | 0.345 |
| condition | 82.67 | 1 | **< 0.001** |
| resident number | 52.03 | 3 | **< 0.001** |
| pyoverdine production | 24.50 | 1 | **< 0.001** |
| habitat : condition | 31.58 | 1 | **< 0.001** |
| habitat : resident number | 2.79 | 3 | 0.425 |
| condition : resident number | 0.24 | 3 | 0.971 |
| habitat : pyoverdine production | 12.73 | 1 | **< 0.001** |
| condition : pyoverdine production | 22.01 | 1 | **< 0.001** |
| resident number : pyoverdine production | 0.93 | 3 | 0.817 |
| habitat : condition : pyoverdine production | 12.62 | 1 | **< 0.001** |

**Figure 4.**
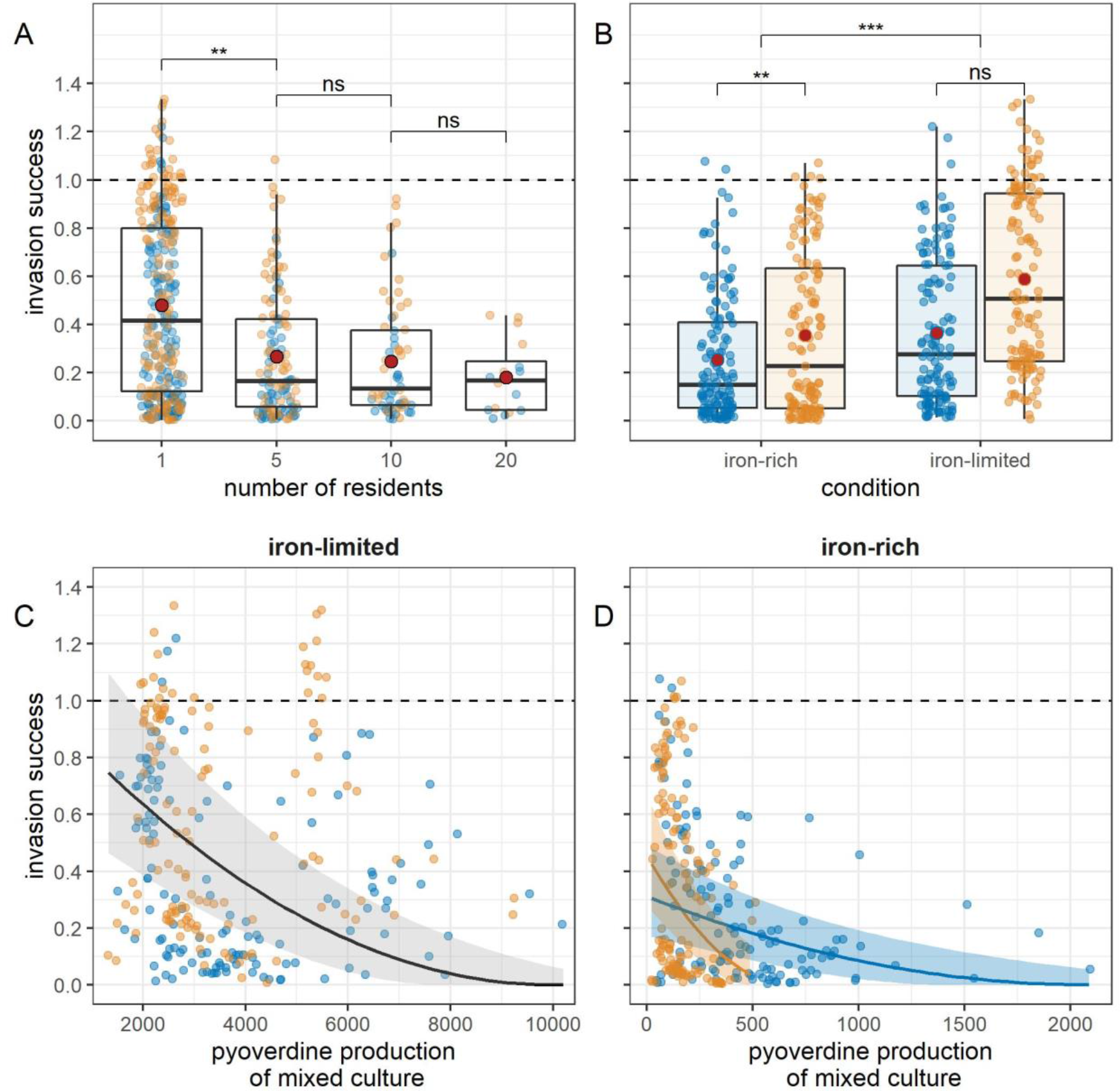
General determinants of the invader’s competitive success. Shown are associations between the invader’s competitive success and (A) the number of soil (orange) and pond (blue) residents, (B) the origin of residents under iron-limited and iron-rich conditions, and the relative pyoverdine production of the mixed culture under (C) iron-limited and (D) iron-rich conditions. Box plots show the median (bold line), mean (red point), 1st and 3rd quartile (box), and 5th and 95th percentile (whiskers). Solid lines and shaded areas are regression lines and 95% confidence intervals. The dashed horizontal lines indicate the null line, where the invader’s growth is not affected by presence of residents (i.e. monoculture growth. The solid and dashed lines in (C) and (D) indicate significant and non-significant relationships, respectively. Shaded areas are 95% confidence intervals (grey = relationship applying to both soil and pond residents; orange = soil-specific relationship; blue = pond-specific relationship).

## Discussion

Although microbial invasions are increasingly receiving attention due to their drastic effects on community composition and functioning, it often remains unclear which microbial traits help invaders to successfully establish themselves in a community (Litchman 2010). Here, we tested whether social interactions mediated by iron-scavenging siderophores predict the success of the opportunistic pathogen *P. aeruginosa* to invade environmental resident communities assembled from 160 *Pseudomonas* isolates from soil and pond samples. We found that pyoverdines – the main siderophores produced by fluorescent pseudomonads – take on a dual role during invasion: they act either as “public goods” by delivering iron to individuals possessing the matching receptors required for uptake, or as “public bads” by locking iron away from competitors lacking these receptors. We show that these pyoverdine-based interactions are a strong predictor of invasion success. Particularly, invasion success generally increased when the residents’ pyoverdines stimulate (public good effect) rather than inhibit (public bad effect) the invader’s growth. This pattern was reinforced in one-to-one competitions when the invader’s own pyoverdine acted as a public bad for the resident. Conversely, invasion success decreased when residents produced inhibitory and/or high amounts of pyoverdines, and when they occurred in more diverse communities.

At the mechanistic level, receptor (in)compatibility should determine whether a pyoverdine acts as a public good or public bad. We previously investigated this aspect through the analysis of 24 resident genomes (Butaite *et al*. 2017). We found that residents had on average 5.1 (range: 1-19) different pyoverdine receptor homologues. Moreover, we found that residents typically produce structurally different pyoverdine types and that the degree of receptor compatibility between residents predicted the level of stimulation versus inhibition (Butaite *et al*. 2017). Given this rich receptor repertoire, it is conceivable that many residents can exploit the invader’s pyoverdine and use it as a growth-stimulating public good in our current setup. This interpretation is further corroborated by our observation that pond residents have significantly more pyoverdine receptor variants than soil residents (median 5.5 vs. 2.5) (Butaite *et al*. 2020), and are significantly better at exploiting the invader’s pyoverdine (Fig. 2B). The situation looks different for our invader *P. aeruginosa* PAO1, which has only two pyoverdine receptors (Ghysels *et al*. 2004). This limited receptor repertoire could explain why most resident pyoverdines had an inhibitory public bad effect on the invader.

In addition to the effect of receptor compatibility, we found that invasion success declined with higher community-wide pyoverdine production levels. This finding indicates that residents with high pyoverdine production levels are better at resisting invasion. This result makes intuitive sense because the pyoverdines of residents are predominantly public bads for the invader, such that higher production levels should lead to higher inhibition. To confirm this conclusion, we ran additional analyses linking the pyoverdine production and growth of residents in monoculture to the effect of their pyoverdine on invader growth (see the Supplementary Material). These analyses confirmed that residents that grow well and produce high pyoverdine amounts are better at inhibiting the invader (Fig. S3).

Since both public bad effects and pyoverdine production levels seem to drive invasion success, we further examined which of the two effects is more important. We thus repeated our analyses of invasion success using a pyoverdine effect that is corrected for the amount of pyoverdine produced by residents (Fig. S4). We found that the corrected pyoverdine effects still positively affected invasion success in competitions with soil residents, but no longer in competitions with pond residents. This suggest that net public bad effects play a more important role in soil communities, whereas the amount of pyoverdine is more relevant in pond communities. This latter finding could be a consequences of our observation that pond residents produce more pyoverdine than soil residents (Fig. 1B), presumably because iron is more limited in the pond habitat (Butaite et al., 2018). Moreover, it could explain why invasion success was lower in pond as compared to soil communities under iron-limited conditions.

Our findings highlight the role of siderophores in driving invasion patterns, but how do they relate to other factors determining invasion outcomes? Consistent with previous work (Bonanomi *et al*. 2014; Acosta *et al*. 2015), we found that invasion success decreased with higher resident numbers. Under iron-limited conditions, this effect presumably arises because the likelihood of public-bad producing residents being present increases with resident number. Under iron-rich conditions, this explanation does not hold, as pyoverdines are not required for growth and therefore not produced in high amounts. Here, we propose that intense resource and interference competition could jointly contribute to this effect. The CAA medium used is a protein digest consisting of amino acids and peptides. While competition under iron-limitation solely revolves around iron, CAA supplemented with iron offers many different nutrient niches. Niche occupancy likely increases with resident number, thereby reducing open niche space for invaders. Although our supernatant assays revealed little evidence for inhibitory effects, invasion success in more diverse communities might have been further reduced by increased interference competition under iron-rich conditions. This is because interference traits such as toxins and contact-dependent killing systems are often only deployed when sensing competitors, especially at high cell density as achieved in iron-rich environments (Melvin *et al*. 2017; Mavridou *et al*. 2018). A combination of intense nutrient and interference competition could finally also explain why invasion success was overall lower under iron-rich than iron-limited conditions.

In the introduction, we proposed that siderophores could mediate both resource and interference competition through their dual role as public good vs. public bad. The resource competition side is obvious, as siderophores make iron available for close relatives (public good effect) but withhold it from competitors lacking a matching receptor (public bad effect). The interference competition side is less established. Although we observed strong inhibition, public bad effects could simply be a by-product of the producer’s iron acquisition strategy. The question is whether producers are selected to secrete siderophores beyond of what is required for iron scavenging to specifically combat competitors. A recent theoretical study showed that siderophore secretion can indeed evolve to inhibit other strains under a wide range of conditions (Niehus *et al*. 2017). Moreover, bacteria have been observed to upregulate siderophore production in the presence of competing species (Harrison *et al*. 2008; Leinweber *et al*. 2018). To elucidate the role of siderophores in interference competition, future studies should focus on the fitness consequences of such siderophore expression adjustments.

While we focused on pyoverdines in *Pseudomonas* communities, a key open question is whether our findings can be extrapolated to diverse microbial consortia with more diverse siderophore and receptor repertoires. A recent study indeed supports such a notion. Gu et al (2020) competed 2150 bacterial isolates from rhizosphere microbiomes against the plant pathogen *Ralstonia solanacearum*. They found that the effects of rhizosphere siderophores on pathogen growth vary from strong growth inhibition to promotion. While no community effects were examined and the type of siderophores and receptors produced remained unknown, this study supports the dual role of siderophores in driving social interactions not only among pseudomonads, but also in diverse microbial communities.

In conclusion, we show that the dual role of siderophores as public good vs. public bad predicts microbial invasions under iron limitation, a condition that prevails in many natural habitats (Boyd & Ellwood 2010; Colombo *et al*. 2014). Our results could have broad implications on two fronts. From a medical perspective, they suggest that siderophore and receptor repertoires of pathogens (such as *P. aeruginosa*) affect their potential to establish reservoirs in natural communities after escaping from patients or hospitals (Loveday *et al*. 2014; Quick *et al*. 2014). From an ecological disease management perspective, our results highlight that knowledge on siderophore and receptor repertoires could be leveraged to engineer microbial communities that protect plant and animal hosts by preventing pathogen invasions through their non-exploitable and pathogen-inhibiting siderophores.

## Acknowledgements

We thank Elena Butaitė for collecting the natural isolates, and Felix Moerman, Zhong Wei, and Andreas Wagner for comments. This research was supported by the German Science Foundation (DFG; KR 5017/2-1 to JK), the University Research Priority Program (URPP) “Evolution in Action” of the University of Zurich, the Swiss National Science Foundation (31003A_182499 to RK), and the European Research Council (ERC) under the European Union’s Horizon 2020 research and innovation programme (grant agreement no. 681295 to RK).

## References

Acosta, F., Zamor, R.M., Najar, F.Z., Roe, B.A. & Hambright, K.D. (2015). Dynamics of an experimental microbial invasion. Proc. Natl. Acad. Sci. U. S. A., 112, 11594–11599.

Amalfitano, S., Coci, M., Corno, G. & Luna, G.M. (2015). A microbial perspective on biological invasions in aquatic ecosystems. Hydrobiologia, 746, 13–22.

Andrews, S.C., Robinson, A.K. & Rodríguez-Quiñones, F. (2003). Bacterial iron homeostasis. FEMS Microbiol. Rev., 27, 215–237.

Bäckhed, F., Ley, R.E., Sonnenburg, J.L., Peterson, D.A. & Gordon, J.I. (2005). Host-bacterial mutualism in the human intestine. Science., 307, 1915–1920.

Bates, D., Mächler, M., Bolker, B.M. & Walker, S.C. (2015). Fitting linear mixed-effects models using lme4. J. Stat. Softw., 67, 1–48.

Bäumler, A.J. & Sperandio, V. (2016). Interactions between the microbiota and pathogenic bacteria in the gut. Nature, 535, 85–93.

Bell, T., Lilley, A.K., Hector, A., Schmid, B., King, L. & Newman, J.A. (2009). A linear model method for biodiversity–ecosystem functioning experiments. Am. Nat., 174, 836–849.

Bonanomi, G., Capodilupo, M., Incerti, G., Gaglione, S.A. & Scala, F. (2014). Fungal diversity increases soil fungistasis and resistance to microbial invasion by a non resident species. Biol. Control, 72, 38–45.

Boyd, P.W. & Ellwood, M.J. (2010). The biogeochemical cycle of iron in the ocean. Nat. Geosci., 3, 675–682.

Brown, S.P., Le Chat, L., De Paepe, M. & Taddei, F. (2006). Ecology of microbial invasions: amplification allows virus carriers to invade more rapidly when rare. Curr. Biol., 16, 2048–2052.

Bruce, J.B., Cooper, G.A., Chabas, H., West, S.A. & Griffin, A.S. (2017). Cheating and resistance to cheating in natural populations of the bacterium Pseudomonas fluorescens. Evolution, 71, 2484–2495.

Butaite, E., Baumgartner, M., Wyder, S. & Kümmerli, R. (2017). Siderophore cheating and cheating resistance shape competition for iron in soil and freshwater Pseudomonas communities. Nat. Commun., 8, 1–12.

Butaite, E., Kramer, J. & Kümmerli, R. (2020). Local adaptation, geographical distance and phylogenetic relatedness: assessing the drivers of siderophore-mediated social interactions in natural bacterial communities. bioRxiv.

Butaite, E., Kramer, J., Wyder, S. & Kümmerli, R. (2018). Environmental determinants of pyoverdine production, exploitation and competition in natural Pseudomonas communities. Environ. Microbiol., 20, 3629–3642.

Colombo, C., Palumbo, G., He, J.Z., Pinton, R. & Cesco, S. (2014). Review on iron availability in soil: interaction of Fe minerals, plants, and microbes. J. Soils Sediments, 14, 538–548.

Cornelis, P. (2010). Iron uptake and metabolism in pseudomonads. Appl. Microbiol. Biot., 86, 1637–1645.

Cornelis, P. & Matthijs, S. (2002). Diversity of siderophore-mediated iron uptake systems in fluorescent pseudomonads: not only pyoverdines. Environ. Microbiol., 4, 787–798.

Doherty, T.S., Glen, A.S., Nimmo, D.G., Ritchie, E.G. & Dickman, C.R. (2016). Invasive predators and global biodiversity loss. Proc. Natl. Acad. Sci. U. S. A., 113, 11261–11265.

Ehrenfeld, J.G. (2010). Ecosystem consequences of biological invasions. Annu. Rev. Ecol. Evol. Syst., 41, 59–80.

Eisenhauer, N., Schulz, W., Scheu, S. & Jousset, A. (2013). Niche dimensionality links biodiversity and invasibility of microbial communities. Funct. Ecol., 27, 282–288.

Van Elsas, J.D., Chiurazzi, M., Mallon, C.A., Elhottovaa, D., Krištufek, V. & Salles, J.F. (2012). Microbial diversity determines the invasion of soil by a bacterial pathogen. Proc. Natl. Acad. Sci. U. S. A., 109, 1159–1164.

Fox, J. & Weisberg, S. (2019). An R Companion to Applied Regression. Third edition. Sage, Thousand Oaks, CA.

García-Bayona, L. & Comstock, L.E. (2018). Bacterial antagonism in host-associated microbial communities. Science, 361, 1–11.

Ghysels, B., Dieu, B.T.M., Beatson, S.A., Pirnay, J.P., Ochsner, U.A., Vasil, M.L., et al. (2004). FpvB, an alternative type I ferripyoverdine receptor of Pseudomonas aeruginosa. Microbiology, 150, 1671–1680.

Gu, S., Wei, Z., Shao, Z., Friman, V.P., Cao, K., Yang, T., et al. (2020). Competition for iron drives phytopathogen control by natural rhizosphere microbiomes. Nat. Microbiol., 5, 1002–1010.

Harrison, F., Paul, J., Massey, R.C. & Buckling, A. (2008). Interspecific competition and siderophore-mediated cooperation in Pseudomonas aeruginosa. ISME J., 2, 49–55.

Hartney, S.L., Mazurier, S., Kidarsa, T.A., Quecine, M.C., Lemanceau, P. & Loper, J.E. (2011). TonB-dependent outer-membrane proteins and siderophore utilization in Pseudomonas fluorescens Pf-5. BioMetals, 24, 193–213.

Kramer, J., López Carrasco, M.Á. & Kümmerli, R. (2020a). Positive linkage between bacterial social traits reveals that homogeneous rather than specialised behavioral repertoires prevail in natural Pseudomonas communities. FEMS Microbiol. Ecol., 96, 1–10.

Kramer, J., Özkaya, Ö. & Kümmerli, R. (2020b). Bacterial siderophores in community and host interactions. Nat. Rev. Microbiol., 18, 152–163.

Kümmerli, R., Schiessl, K.T., Waldvogel, T., Mcneill, K. & Ackermann, M. (2014). Habitat structure and the evolution of diffusible siderophores in bacteria. Ecol. Lett., 17, 1536– 1544.

Kuznetsova, A., Brockhoff, P.B. & Christensen, R.H.B. (2017). lmerTest package: tests in linear mixed effects models. J. Stat. Softw., 82, 1.26.

Leinweber, A., Weigert, M. & Kümmerli, R. (2018). The bacterium Pseudomonas aeruginosa senses and gradually responds to interspecific competition for iron. Evolution, 72, 1515–1528.

Li, S., Tan, J., Yang, X., Ma, C. & Jiang, L. (2019). Niche and fitness differences determine invasion success and impact in laboratory bacterial communities. ISME J., 13, 402–412.

Libberton, B., Horsburgh, M.J. & Brockhurst, M.A. (2015). The effects of spatial structure, frequency dependence and resistance evolution on the dynamics of toxin-mediated microbial invasions. Evol. Appl., 8, 738–750.

Litchman, E. (2010). Invisible invaders: non-pathogenic invasive microbes in aquatic and terrestrial ecosystems. Ecol. Lett., 13, 1560–1572.

Loveday, H.P., Wilson, J.A., Kerr, K., Pitchers, R., Walker, J.T. & Browne, J. (2014). Association between healthcare water systems and Pseudomonas aeruginosa infections: a rapid systematic review. J. Hosp. Infect., 86, 7–15.

Lowe, S., Browne, M., Boudjelas, S. & De Poorter, M. (2000). 100 of the world’s worst invasive alien species: a selection from the global invasive species database. Invasive Species Specialist Group., Auckland.

Ma, C., Liu, M., Wang, H., Chen, C., Fan, W., Griffiths, B., et al. (2015). Resource utilization capability of bacteria predicts their invasion potential in soil. Soil Biol. Biochem., 81, 287–290.

Mächler, E. & Altermatt, F. (2012). Interaction of species traits and environmental disturbance predicts invasion success of aquatic microorganisms. PLOS ONE, 7, 1–10.

Mallon, C.A., Van Elsas, J.D. & Salles, J.F. (2015a). Microbial invasions: the process, patterns, and mechanisms. Trends Microbiol., 23, 719–729.

Mallon, C.A., Poly, F., Le Roux, X., Marring, I., Van Elsas, J.D. & Salles, J.F. (2015b). Resource pulses can alleviate the biodiversity-invasion relationship in soil microbial communities. Ecology, 96, 915–926.

Mangiafico, S. (2018). rcompanion: functions to support extension education program evaluation. Available at: https://cran.r-project.org/web/packages/rcompanion/index.html. Last accessed, 02/05/2021

Mavridou, D.A.I., Gonzalez, D., Kim, W., West, S.A. & Foster, K.R. (2018). Bacteria use collective behavior to generate diverse combat strategies. Curr. Biol., 28, 345–355.

Melvin, J.A., Gaston, J.R., Phillips, S.N., Springer, M.J., Marshall, C.W., Shanks, R.M.Q., et al. (2017). Pseudomonas aeruginosa contact-dependent growth inhibition plays dual role in host-pathogen interactions. mSphere, 2, 1–13.

Meyer, J.M., Gruffaz, C., Raharinosy, V., Bezverbnaya, I., Schäfer, M. & Budzikiewicz, H. (2008). Siderotyping of fluorescent Pseudomonas: molecular mass determination by mass spectrometry as a powerful pyoverdine siderotyping method. BioMetals, 21, 259– 271.

Van Nevel, S., De Roy, K. & Boon, N. (2013). Bacterial invasion potential in water is determined by nutrient availability and the indigenous community. FEMS Microbiol. Ecol., 85, 593–603.

Niehus, R., Picot, A., Oliveira, N.M., Mitri, S. & Foster, K.R. (2017). The evolution of siderophore production as a competitive trait. Evolution, 71, 1443–1455.

Quick, J., Cumley, N., Wearn, C.M., Niebel, M., Constantinidou, C., Thomas, C.M., et al. (2014). Seeking the source of Pseudomonas aeruginosa infections in a recently opened hospital: an observational study using whole-genome sequencing. BMJ Open, 4, 1–10.

Rezzoagli, C., Granato, E.T. & Kümmerli, R. (2019). In-vivo microscopy reveals the impact of Pseudomonas aeruginosa social interactions on host colonization. ISME J., 13, 2403– 2414.

Schalk, I.J., Rigouin, C. & Godet, J. (2020). An overview of siderophore biosynthesis among fluorescent Pseudomonads and new insights into their complex cellular organization. Environ. Microbiol., 22, 1447–1466.

Sexton, D.J., Glover, R.C., Loper, J.E. & Schuster, M. (2017). Pseudomonas protegens Pf-5 favours self-produced siderophore over free-loading in interspecies competition for iron. Environ. Microbiol., 19, 3514–3525.

Stover, C.K., Pham, X.Q., Erwin, A.L., Mizoguchi, S.D., Warrener, P., Hickey, M.J., et al. (2000). Complete genome sequence of Pseudomonas aeruginosa PAO1, an opportunistic pathogen. Nature, 406, 959–964.

Sunagawa, S., Coelho, L.P., Chaffron, S., Kultima, J.R., Labadie, K., Salazar, G., et al. (2015). Structure and function of the global ocean microbiome. Science, 348, 1–10.

Thakur, M.P., van der Putten, W.H., Cobben, M.M.P., van Kleunen, M. & Geisen, S. (2019). Microbial invasions in terrestrial ecosystems. Nat. Rev. Microbiol., 17, 621–631.

Trivedi, P., Leach, J.E., Tringe, S.G., Sa, T. & Singh, B.K. (2020). Plant–microbiome interactions: from community assembly to plant health. Nat. Rev. Microbiol., 18, 607– 621.

Wagg, C., Bender, S.F., Widmer, F. & Van Der Heijden, M.G.A. (2014). Soil biodiversity and soil community composition determine ecosystem multifunctionality. Proc. Natl. Acad. Sci. U. S. A., 111, 5266–5270.

Wei, Z., Yang, T., Friman, V.P., Xu, Y., Shen, Q. & Jousset, A. (2015). Trophic network architecture of root-associated bacterial communities determines pathogen invasion and plant health. Nat. Commun., 6, 1–9.

Yang, T., Wei, Z., Friman, V.P., Xu, Y., Shen, Q., Kowalchuk, G.A., et al. (2017). Resource availability modulates biodiversity-invasion relationships by altering competitive interactions. Environ. Microbiol., 19, 2984–2991.

