## Supplementary Material for "Siderophores drive invasion dynamics in bacterial communities through their dual role as public good versus public bad"

This file contains the following Supplementary Information:

- **Supplementary Analyses**
- **Supplementary Table S1+S2**
- **Supplementary Figure S1-S4**

**Supplementary Analyses**

**Pyoverdine vs. overall siderophore production.** The net growth effect of the residents' pyoverdines [$GE_{net}$] could theoretically also reflect the action of siderophores other than pyoverdine. To account for this possibility, we measure the total amount of siderophores in the supernatant generated by our residents under iron-limited conditions using the Chrome Azurol Sulfonate (CAS) assay (Schwyn and Neilands 1987; DOI: 10.1016/0003-2697(87)90612-9). This colorimetric assay utilizes an Fe(III)-Chrome Azurol Sulfonate-Hexadecyltrimethylammonium bromide [Fe(III)-CAS-HDTMA] complex that changes color, from blue to orange, when it is deferrated by exchange reactions with siderophores. We first mixed the CAS-solution according to the original recipe (see Schwyn and Neilands 1987 for details) and stored it in the dark until use. To measure the total siderophore amount, we transferred 20 µL of each supernatant into plates filled with 240 µL of CAS assay solution per well. The inoculated plates were incubated at room temperature in the dark for 30 minutes. After that, we quantified the residents' overall production of siderophores by measuring the loss of the blue-colored Fe-CAS-HDTMA complex at 630 nm.

To examine the association between the residents' pyoverdine production (measured based on its natural fluorescence; see the main text) and their overall production of siderophores, we calculated a linear mixed model using the (square root-transformed inverse of the) amount of siderophores in the supernatant as a response variable, and pyoverdine production, resident habitat (soil or pond), as well as their interaction as explanatory variables. Resident community was fitted as a random effect to account for the non-independence of measurements of residents from the same community. Model selection was performed as described in the main text. We found a strong positive association between the residents' production of siderophores and their pyoverdine production ($t_{156} = 14.97$, $p < 0.001$; Fig. S2). By contrast, siderophore production did not differ between habitats (main effect: $F_1 = 2.76$, $p = 0.097$; interaction with pyoverdine production: $F_1 = 2.02$, $p = 0.157$ before removal of the interaction from the model).

**The effect of resident pyoverdines depends on resident growth.** In the main text, we show that invasion success was negatively correlated with the overall pyoverdine production of the mixed culture. This suggests that the effect of resident pyoverdines on invader growth might not only depend on whether or not the invader has a matching receptor for pyoverdine uptake, but also on the amount of pyoverdine produced by the focal resident(s). To explore this possibility, we corrected the effect of resident pyoverdines for differences in pyoverdine amount, and then re-ran our analyses of invasion success with these 'corrected' pyoverdine effects. In the first step, we fitted a linear model using the (square root-transformed) effect of the residents' pyoverdines as a response, and their pyoverdine production as explanatory variable. As the plot of the raw data suggested a non-linear relationship, we fitted the model with a quadratic term. Our model indeed revealed a non-linear decrease of the effect of the residents' pyoverdines with their pyoverdine production levels (linear term: $t_{156} = -10.961$, $p < 0.001$; quadratic term: $t_{156} = -3.488$, $p < 0.001$; Fig. S3A). To obtain a corrected pyoverdine effect of single residents, we extracted the residuals from this model. To be able to link invasion success to these corrected pyoverdine effects in multi-resident mixes, we then calculated the mean corrected value across all residents in a mix. Finally, note that the pyoverdine effect is not only correlated with the amount of pyoverdine, but also with the growth of the pyoverdine-producing culture (linear term: $t_{157} = - 9.204$, $p < 0.001$; quadratic term: $t_{156} = 1.357$, $p = 0.177$ before removal of the term; Fig. S3B). This is expected, as growth and pyoverdine production are themselves linked (see the main text). We corrected for differences in pyoverdine production instead of differences in growth, as it seems more apt to assume that pyoverdine production allows for growth (rather than the other way round) under iron-limitation, our condition of interest.

In the second step, we re-ran our analyses of invasion success using the corrected pyoverdine effects. In contrast to the analyses presented in the main text, invasion success between the invader and single residents was no longer shaped by an interaction between the invader's and the (now corrected) resident's siderophore effects ($\chi^2_1 = 0.518$, $p = 0.472$ before removal of the interaction from the model).

Instead, invasion success was overall negatively correlated with the effect of the invader's pyoverdine on the residents ($t_{151.22} = -2.172$, $p = 0.031$), whereas the corrected effect of the resident's pyoverdine on the invader depended on resident habitat (corrected siderophore effect: $\chi^2_1 = 20.979$, $p < 0.001$; habitat: $\chi^2_1 = 2.759$, $p = 0.097$; interaction: $\chi^2_1 = 8.901$, $p = 0.003$). Specifically, invasion success only increased with the corrected effect of the residents' pyoverdines in soil residents ($t_{152.41} = 5.453$, $p < 0.001$), but not in pond residents ($t_{148.69} = 0.371$, $p = 0.711$; Figure S4A). Interestingly, we found the same interaction to shape invasion success in competitions with multiple residents (Table S2), i.e. the corrected effect of the residents' pyoverdines was positively correlated with invasion success in communities of soil ($t_{95.09} = 2.752$, $p = 0.007$), but not pond residents ($t_{93.69} = -1.328$, $p = 0.187$; Figure S4B).

Overall, these analyses show that both receptor compatibility and the amount of pyoverdine jointly contribute to the effect of pyoverdine on invader growth. The fact that the corrected effect of the residents' siderophores no longer influenced invasion success in competitions with pond residents, whereas its influence persisted in competitions with soil residents, shows that habitat-specific differences in the type and amount of pyoverdines can affect invasion outcomes. Intriguingly, it also suggests that the degree of receptor compatibility can vary. In soil residents, the effect of resident pyoverdines on invasion success persisted even after accounting for differences in production levels. This suggests the invader might still be able to import some resident pyoverdines at low rate (for instance if they are structurally similar to its own), which would diminish the negative effect arising from their locking away of iron and might hence increase the invader's success. In competitions with pond residents, invasion success was affected by the overall effect of resident pyoverdines, but not by the 'amount-corrected' effect. This suggests that there might be little structural variation among the pyoverdines of pond residents, such that the overall effect of these pyoverdines is mostly driven by differences in pyoverdine amount.

**Table S1 | Predictors of resident growth.** Significant p-values in bold.

| | $\chi^2_1$ | p |
|---|---|---|
| condition | 804.26 | **< 0.001** |
| habitat | 0.18 | 0.673 |
| pyoverdine production | 199.58 | **< 0.001** |
| condition : habitat | 0.50 | 0.480 |
| condition : pyoverdine production | 28.12 | **< 0.001** |
| habitat : pyoverdine production | 1.02 | 0.313 |
| condition : habitat : pyoverdine production | 21.10 | **< 0.001** |

**Table S2 | Predictors of invasion success into multi-resident communities.** Significant p-values in bold.

| | $\chi^2_1$ | p |
|---|---|---|
| habitat | 5.43 | **0.020** |
| resident number | 1.55 | 0.213 |
| effect of invader siderophores on resident growth (IoR) | 1.15 | 0.284 |
| corrected effect of resident siderophores on invader (RoI) | 1.26 | 0.262 |
| habitat : RoI | 7.96 | **0.005** |

**Resident (R)**

| **Invader (I)** | Public bad | Pulic good |
|---|---|---|
| Public bad | Conditional | Invasion |
| Public good | Exclusion | Conditional |

**Figure S1 | Conceptual 2x2 interaction matrix of expected invasion outcomes.** The matrix depicts the expected invasion outcomes based on whether a focal resident and the invader produce pyoverdines that act as public good or public bad, respectively. If the invader produces a public good and the resident a public bad, invasion is unlikely to be successful because the resident can both exploit the invader's pyoverdine and use its own pyoverdine to lock iron away. Conversely, if the invader produces a public bad and the resident a public good, invasion is expected to be successful, as the invader can exploit the resident's pyoverdine and use its own pyoverdine to lock iron away. Finally, if both the resident and the invader produce public goods or public bads, invasion success will be conditional on other factors, such as the amount of secreted pyoverdines and the magnitude of their inhibitory or stimulatory effects.

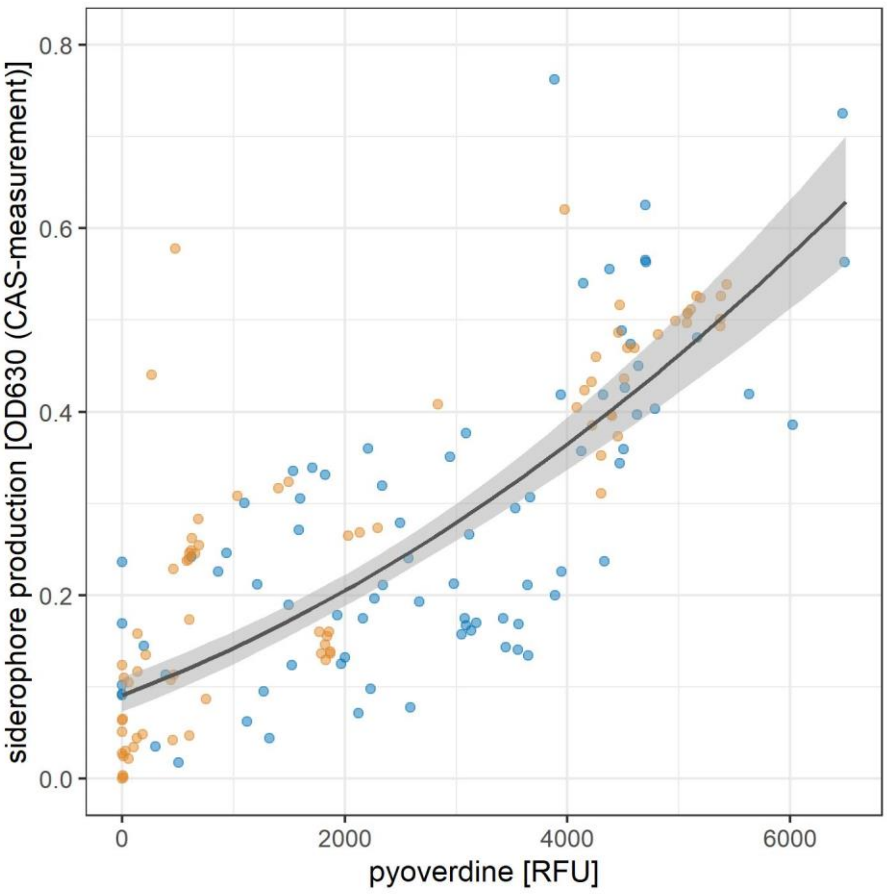


**Figure S2 | Positive association between the residents' pyoverdine and overall siderophore production.** Shown is the (habitat-independent) increase of the overall siderophore production of pond (blue) and soil (orange) residents under iron-limited conditions with the production of pyoverdine, the main type of siderophore produced by fluorescent pseudomonads. The grey-shaded area is the 95% confidence interval.

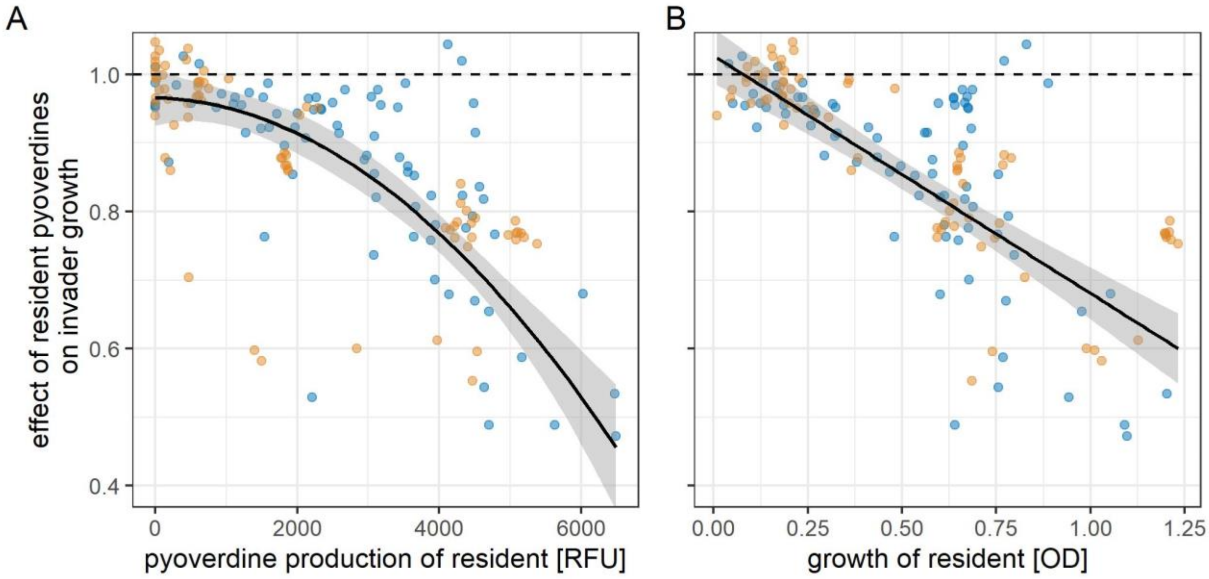


**Figure S3 | Decrease of the effect of resident pyoverdines with resident pyoverdine production and growth.** Shown is the decrease of the effect of resident pyoverdines on invader growth with (A) the pyoverdine production and (B) the growth of pond (blue) and soil (orange) residents. Shaded areas are 95% confidence intervals (grey = relationship applying to both soil and pond residents). The dashed horizontal line is a null line indicating no effect of a resident's pyoverdine on invader growth.

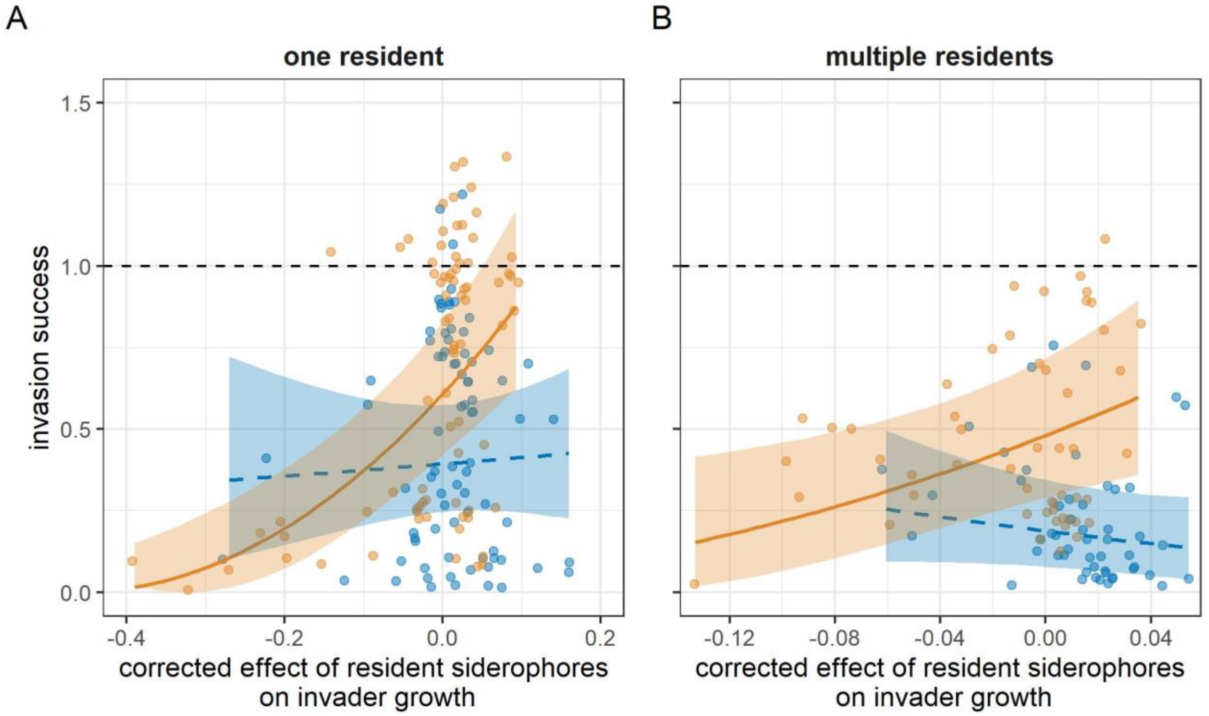


**Figure S4 | Invasion success and the corrected pyoverdine effect.** Depicted are the links between invasion success and the effect of resident siderophores on invader growth (after correction for differences in pyoverdine production among residents) in competitions between the invader and (A) one resident or (B) multiple residents from pond (blue) and soil (orange) communities. Shaded areas are 95% confidence intervals (grey = relationship applying to both soil and pond residents; orange = soil-specific relationship; blue = pond-specific relationship). Solid and dashed regression lines indicate significant and non-significant relationships, respectively. The dashed horizontal lines are null lines indicating no effect of resident pyoverdine(s) on invader growth.
